# Quantifying compositional variability in microbial communities with FAVA

**DOI:** 10.1101/2024.07.03.601929

**Authors:** Maike L. Morrison, Katherine S. Xue, Noah A. Rosenberg

## Abstract

Microbial communities vary across space, time, and individual hosts, presenting new challenges for the development of statistics measuring the variability of community composition. To understand differences across microbiome samples from different host individuals, sampling times, spatial locations, or experimental replicates, we present FAVA, a new normalized measure for characterizing compositional variability across multiple microbiome samples. FAVA quantifies variability across many samples of taxonomic or functional relative abundances in a single index ranging between 0 and 1, equaling 0 when all samples are identical and equaling 1 when each sample is entirely comprised of a single taxon. Its definition relies on the population-genetic statistic *F*_*ST*_, with samples playing the role of “populations” and taxa playing the role of “alleles.” Its convenient mathematical properties allow users to compare disparate data sets. For example, FAVA values are commensurable across different numbers of taxonomic categories and different numbers of samples considered. We introduce extensions that incorporate phylogenetic similarity among taxa and spatial or temporal distances between samples. We illustrate how FAVA can be used to describe across-individual taxonomic variability in ruminant microbiomes at different regions along the gastrointestinal tract. In a second example, a longitudinal analysis of gut microbiomes of healthy human adults taking an antibiotic, we use FAVA to quantify the increase in temporal variability of microbiomes following the antibiotic course and to measure the duration of the antibiotic’s influence on microbial variability. We have implemented this tool in an R package, *FAVA*, which can fit easily into existing pipelines for the analysis of microbial relative abundances.

**Significance statement:** Studies of microbial community composition across time, space, or biological replicates often rely on summary statistics that analyze just one or two samples at a time. Although these statistics effectively summarize the diversity of one sample or the compositional dissimilarity between two samples, they are ill-suited for measuring variability across many samples at once. Measuring compositional variability among many samples is key to understanding the temporal stability of a community across multiple time points, or the heterogeneity of microbiome composition across multiple experimental replicates or host individuals. Our proposed measure, FAVA, meets the need for a statistic summarizing compositional variability across many microbiome samples all at once.

## Introduction

Understanding the compositional variability of microbial communities across space, time, or host individuals is important for characterizing these communities and their relationships with biological variables of interest (Turnbaugh *et al*., 2007; Dethlefsen and Relman, 2011; Faith *et al*., 2013; David *et al*., 2014; Flores *et al*., 2014; Coyte *et al*., 2015; Oh *et al*., 2016; Thompson *et al*., 2017; Goldford *et al*., 2018; Ji *et al*., 2019; Fassarella *et al*., 2021; Estrela *et al*., 2022; Upadhyay *et al*., 2023). For example, studies of microbiome composition have found that microbiome compositions are often more variable across dysbiotic individuals than across healthy individuals (Zaneveld *et al*., 2017), the microbial communities of infants tend to be more variable across individuals than those of adults (Kurokawa *et al*., 2007), and gut and tongue microbiomes that are more diverse may be less temporally variable (Flores *et al*., 2014). Despite its biological importance, however, compositional variability is difficult to directly quantify with existing methods. We define “compositional variability” as variability across two or more compositional vectors— lists of proportions that sum to 1 (Figure 1A). Compositional variability is minimized when the compositional vectors have identical compositions; it is maximized when each vector contains a single category at 100% frequency (Figure 1B and C). We focus on vectors that represent the composition of microbiome samples. These vectors’ entries represent relative abundances of taxonomic categories such as OTUs, species, or even functional categories such as gene classifications, inferred from 16S or metagenomic sequencing data (Lozupone *et al*., 2012; Louca *et al*., 2017; Shalon *et al*., 2023). Each vector can represent the composition of a microbiome sample from a distinct timepoint, spatial location, host individual, or replicate. Compositional variability can therefore represent temporal stability, spatial heterogeneity, inter-host diversity, or repeatability (Faith *et al*., 2013; Bashan *et al*., 2016; Goldford *et al*., 2018; Mehta *et al*., 2018; Seekatz *et al*., 2019; Sheth *et al*., 2019; Roodgar *et al*., 2021; Estrela *et al*., 2022; Guthrie *et al*., 2022; Shalon *et al*., 2023)

**Figure 1:**
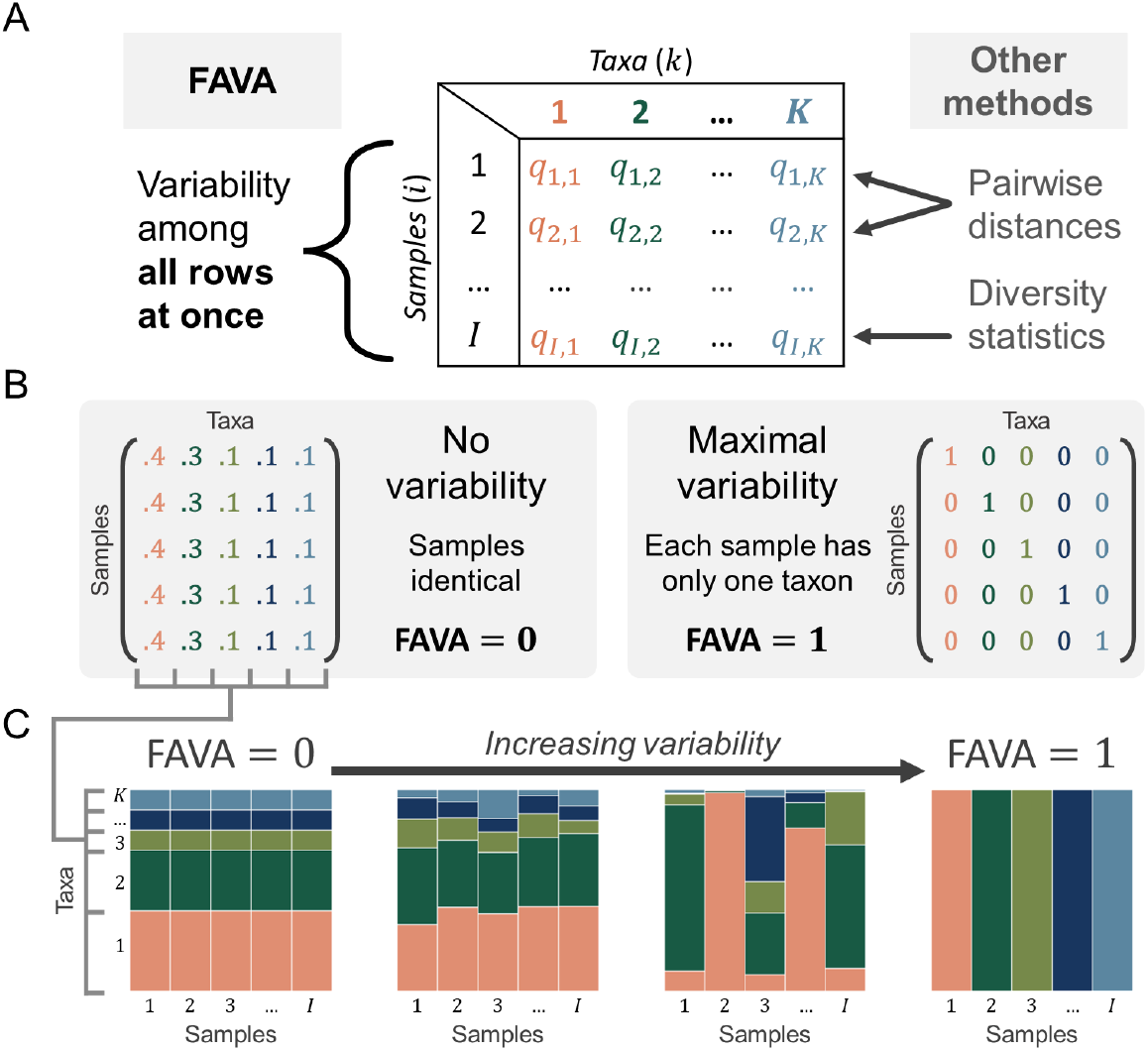
FAVA quantifies compositional variability across many abundance vectors in a single number. **(A)** FAVA is computed among the rows of a matrix, with rows representing microbiome samples, columns corresponding to microbial taxa such as OTUs or families, and entries representing relative abundances. FAVA quantifies variability across many rows in a single number, distinguishing it from other methods, which analyze just one or two rows at a time. **(B)** Given the number of samples (*I*) and number of taxa (*K*), variability is minimized if the rows are identical, corresponding to the case where each sample is the same as every other sample. Variability is maximized if each sample contains just one taxon, as long as there are at least two different taxa present across all samples. When plotted as a relative abundance plot (e.g., panel C), each row of these matrices is visualized as a vertical bar. The matrix pivots 90 degrees when visualized. **(C)** For each matrix, variability across samples increases as the samples become less similar (from left to right). From left to right, the values of FAVA for the matrices are 0, 0.006, 0.452, and 1.

Traditionally, microbiome studies have used statistics such as the Shannon and Gini-Simpson indices (Patil and Taillie, 1982), the Jensen-Shannon divergence (Lin, 1991), and the Bray-Curtis dissimilarity (Bray and Curtis, 1957). Single-sample diversity statistics such as the Shannon and Gini-Simpson indices quantify the variability of microbiome samples considered individually, answering questions such as “Which of these microbiomes is the most diverse?” Pairwise statistics, such as the Jensen-Shannon divergence, Jaccard index, and Bray-Curtis dissimilarity compare the compositions of two samples, answering questions such as “How does the composition of a perturbed microbial community compare to a pre-perturbation reference sample?” Although these tools are valuable when variability is of interest in one sample or between two samples, they are less well suited to scenarios in which three or more samples are of interest, as they only consider one or two samples at once.

Studies that seek to quantify variability across many samples are often limited to computing summary measures of each sample, such as diversity indices (Flores *et al*., 2014; Shalon *et al*., 2023), principal component coefficients (Zaneveld *et al*., 2017; Olm *et al*., 2022), or the abundances of individual taxa (Ji *et al*., 2019; Kang *et al*., 2022), and computing the variability across samples of these summary statistics. However, this approach measures the variability of a summary statistic, not the variability of the microbiome composition itself. Because it is possible for very different compositions to produce similar values of a summary statistic, such indirect variability measures potentially obscure large differences among samples. Existing methods are therefore insufficient to, for example, determine if a microbiome is more temporally variable after than before a perturbation, or to identify which of many anatomical regions has the most across-individual microbiome variability. These questions require statistics of compositional variability that are suited to arbitrary numbers of samples.

Consider for illustration the study of Flores et al. (Flores *et al*., 2014), which aimed to compare regions of the body in terms of their temporal variability in microbiome composition. For 85 adults, they profiled the microbiomes of four body habitats weekly for three months. They measured temporal variability by computing diversity statistics such as the Shannon index for each temporal sample, then computing the coefficient of variation of the Shannon index over time for each of the 85 individuals and four body regions. This approach quantifies the variability of the Shannon diversity, not the variability of the microbiome composition itself. Because microbiomes with either no species or many species in common could have identical Shannon indices, this method could assign time series with dramatically different compositional change the same coefficient of variation, obscuring meaningful differences.

In this paper, we present FAVA, a statistical measure that quantifies variability of microbiome composition across many microbiome samples. In a single number, FAVA measures variability of microbial composition across arbitrarily many microbiome samples, providing a summary of large data sets. The measure allows for the optional inclusion of similarities among taxonomic categories (e.g., phylogenetic similarity) as well as for optional non-uniform weighting of samples (e.g., to account for uneven sampling time intervals). FAVA, which stands for an F_ST_-based Assessment of Variability across vectors of relative Abundances, is based on the population-genetic statistic *F*_*ST*_, which is traditionally used to quantify variability across vectors of allele frequencies for multiple populations. FAVA takes values between 0 and 1, equaling 0 when all sampled microbiome compositions are identical, and equaling 1 when each sample contains only a single taxon and at least two distinct taxa are present across samples (Figure 1B and C). Because FAVA is a normalized statistic, it can be used to compare variability among sets of samples with very different numbers of taxa or datasets with very different numbers of samples.

We demonstrate FAVA with two datasets, one containing spatial samples along the gastrointestinal tract of seven species of ruminants, and the other describing time series of microbiome samples from 22 human individuals who experienced an antibiotic perturbation. In the ruminant dataset, we identify substantially higher inter-individual variability in the stomach and small intestine than in the large intestine, supporting the view that substantial microbiome variability is obscured when gastrointestinal communities are sampled through fecal samples alone (Shalon *et al*., 2023). In the human dataset, we show that temporal variability in microbiome composition is significantly elevated following an antibiotic perturbation, and that just half of subjects return to low levels of temporal variability in the 30 days following completion of the antibiotic.

## Results

### Definition of FAVA

The composition of a microbial community is most commonly described in terms of relative abundances of OTUs, species, bacterial families, or other units, including functional units such as gene categories. Matrices of such abundances are central to software widely used for the analysis of microbiome data, such as *Phyloseq* (McMurdie and Holmes, 2013) and *QIIME2* (Bolyen *et al*., 2019). In an “OTU table,” denoted *Q*, each row represents a microbial community sample, each column represents a distinct taxon, and the entry *q*_*i,k*_ represents the relative abundance of taxon *k* in sample *i* (Figure 1A). The samples in an OTU table represent samples of microbial communities that could vary in their sampling location, sampling time, and subject or replicate. Throughout this paper, we use “sample *i*” to refer to the *i*^*th*^ row of the OTU table.

FAVA quantifies variability across the rows of an OTU table (Figure 1A). If the rows represent samples from different time points for one subject, FAVA is a measure of the temporal stability of the community. If the rows represent different sampling locations, FAVA quantifies the spatial heterogeneity of the community. FAVA can be independently computed on disjoint subsets of the rows of an OTU table. For example, to measure the temporal variability in microbiome composition for each of many subjects, the entire matrix would contain many subjects and time points, and matrix subsets containing just one subject and many time points could be separately analyzed. The measure ranges between 0 (no variability) and 1 (maximum variability), and can thus be used to compare the variabilities of multiple sets of samples (Figure 1B and C).

FAVA is based on the population-genetic statistic *F*_*ST*_, which is traditionally used to measure the variability of allele frequencies across populations but can also be applied to other types of compositional data (Jost *et al*., 2010; Morrison *et al*., 2022, 2023). We apply FAVA to microbiome data by analyzing microbial taxon abundances in place of allele frequencies, and microbiome samples in place of populations.

*F*_*ST*_ is defined in terms of the population-genetic statistic heterozygosity, mathematically equivalent to the Gini-Simpson diversity in ecology. For a sample *i* with *k* = 1, 2, …, *K* taxa with abundances *q*_*i,k*_, the Gini-Simpson diversity of the sample is the probability that two random draws from the sample do not belong to the same taxon (Patil and Taillie, 1982):

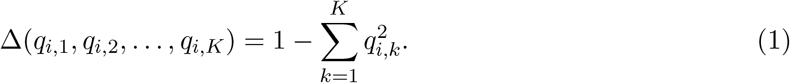

Δ(*q*_*i*,1_, *q*_*i*,2_, …, *q*_*i,K*_) = 0 if and only if some taxon has abundance 1 and all others have abundance 0 (i.e., *q*_*i,k*_*′* = 1 for some *k*^*′*^, and *q*_*i,k*_ = 0 for all *k ≠ k*^*′*^). Δ(*q*_*i*,1_, *q*_*i*,2_, …, *q*_*i,K*_) = 1 − ^1^, its maximum given *K*, if and only if all taxa are equally abundant (i.e., 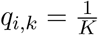 for all *k* = 1, 2, …, *K*).

*F*_*ST*_ proceeds by computing this diversity index on the set of all *i* = 1, 2, …, *I* microbiome samples (i.e., rows of the OTU table, *Q*) in two ways. The mean sample Gini-Simpson diversity, Δ_*S*_, is computed by averaging the Gini-Simpson diversities of the samples:

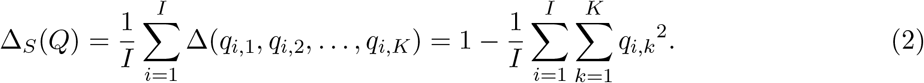

The total Gini-Simpson diversity, Δ_*T*_, is the Gini-Simpson index if the samples were pooled. It is computed by first calculating the centroid of the samples (the vector of mean taxon abundances over all *I* samples) and then computing the Gini-Simpson diversity of the centroid:

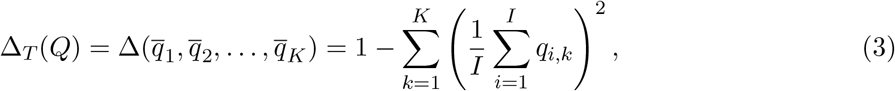

where 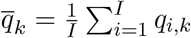. In short, we compute Δ_*S*_ by first computing the Gini-Simpson index for all samples and then averaging, and we compute Δ_*T*_ by first averaging all samples and then computing the Gini-Simpson index.

The population-genetic statistic *F*_*ST*_ is the normalized difference between these two quantities:

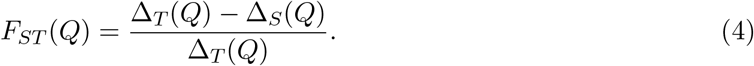

Assuming Δ_*T*_ (*Q*) > 0, *F*_*ST*_ equals 0 if and only if Δ_*T*_ (*Q*) = Δ_*S*_(*Q*), which occurs if and only if all *I* samples are identical (Figure 1B, left-hand side). *F*_*ST*_ equals 1 if and only if Δ_*S*_(*Q*) = 0 and Δ_*T*_ (*Q*) > 0, which occurs if and only if each sample has only a single taxon, and there are at least two distinct taxa present across all samples (Figure 1B, right-hand side). In the language of OTU tables, *F*_*ST*_ equals 0 if and only if all rows of the OTU table are identical, and it equals 1 if and only if each row contains a single one and *K* − 1 zeroes. *F*_*ST*_ can be viewed as a measure of how well-mixed the samples are across a dimension of interest: if all samples are perfectly well-mixed, then their compositions will be identical and *F*_*ST*_ will equal 0.

Possible values of FAVA range between 0 and 1 for any sample size. However, when the number of samples is small, *F*_*ST*_ can be constrained by the mean frequency of the dominant taxon, especially if this frequency is close to 0 or 1 (Alcala and Rosenberg, 2022). Normalizing *F*_*ST*_ by its theoretical upper bound conditional on the number of samples and the mean frequency of the most abundant taxon 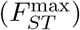 can account for this property, allowing for differences in variability to be distinguished from differences in the abundance of the dominant taxon. However, because the normalized statistic is divided by a theoretical upper bound possibly less than 1, 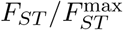 can equal one without satisfying the conditions described in Figure 1C. The normalized statistic 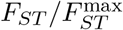 (Morrison *et al*., 2022) is included as an option in the *FAVA* R package. Further discussion of when to consider normalizing *F*_*ST*_ by this upper bound is included in the *FAVA* R package’s vignette on microbiome data analysis.

Having introduced FAVA and established its mathematical properties, we now apply the method to data. Here, we focus on two example applications: using FAVA to quantify variability across individuals in the ruminant gastrointestinal tract, and using FAVA to quantify temporal variability in the human gut in response to antibiotic perturbation.

### Gastrointestinal microbiome variability across ruminant species

Along the vertebrate gastrointestinal tract, factors such as nutrient availability, pH, and oxygen level vary substantially, shaping the types, abundances, and functions of resident microbes (Tropini *et al*., 2017; Folz *et al*., 2023; Shalon *et al*., 2023). Quantifying the across-host variability of microbiomes along the gastrointestinal tract can elucidate spatially-structured, in vivo community assembly.

We here use FAVA to quantify the variability of ruminant gastrointestinal microbiomes across individuals from seven host species. We analyze data from Xie et al. (Xie *et al*., 2021), who used shotgun metagenomics to profile samples collected along the gastrointestinal tracts of 37 individuals across seven species of ruminants (Figure 2). For each individual, Xie et al. (Xie *et al*., 2021) collected samples from ten gastrointestinal regions: the rumen, reticulum, omasum, and abomasum of the stomach (Figure 2, blue *x*-axis labels); the duodenum, jejunum, and ileum of the small intestine (Figure 2, yellow *x*-axis labels); and the cecum, colon, and rectum of the large intestine (Figure 2, red *x*-axis labels). Xie et al. (Xie *et al*., 2021) used their metagenomic sequences to infer abundances of both taxonomic categories, such as microbial genera (Figure 2A), and functional categories, such as carbohydrate-active enzymes (Figure 2C). We computed FAVA on these data in order to understand which gastrointestinal regions are the most and least variable across host individuals (Figure 2C), and to compare the across-host variability of microbial genera to the acrosshost-species variability of functional gene categories throughout the gastrointestinal tract (Figure 2D).

**Figure 2:**
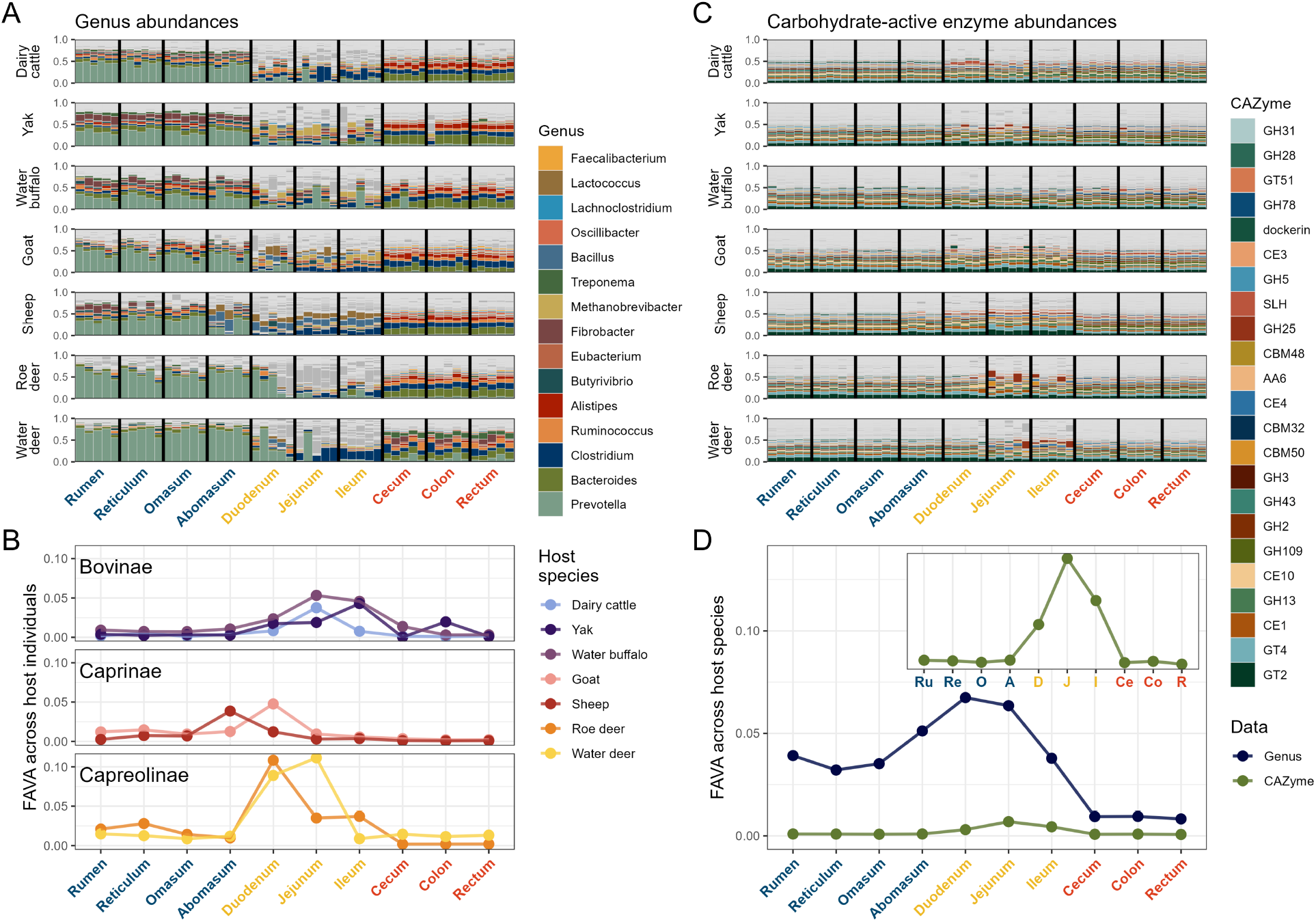
FAVA quantifies taxonomic and functional variability across host individuals and species along the ruminant gastrointestinal tract. **(A) Relative abundances of genera across gastrointestinal regions for seven host species:** dairy cattle (*n* = 6), yak (*n* = 5), water buffalo (*n* = 5), goat (*n* = 6), sheep (*n* = 5), roe deer (*n* = 5), and water deer (*n* = 5). Vertical black lines delimit the 10 gastrointestinal regions: rumen, reticulum, omasum, and abomasum for the stomach (blue); duodenum, jejunum, and ileum for the small intestine (yellow); and cecum, colon, and rectum for the large intestine (red). The ordering of these regions matches the ordering of the regions along the digestive tract. Each colored vertical bar represents the genus-level composition of a microbiome sample from one gastrointestinal region within one host individual. The horizontal ordering of individuals is consistent across regions. Of the 4,134 genera across all samples, the 15 with mean across-sample abundance greater than 1% are colored; all other genera are dark gray. Light gray horizontal lines delineate breaks between different genera. Genera are plotted in order of decreasing genus abundance across all samples and host species, from bottom (most abundant) to top (least abundant). **(B) Across-individual variability of genus abundances within each gastrointestinal region for each host species**. Each dot corresponds to the value of FAVA computed across all samples within a gastrointestinal region for one species. Species are grouped into panels by subfamily. **(C) Carbohydrate-active enzyme (CAZyme) relative abundances across gastrointestinal regions for each host species**. We color the 23 of 350 CAZymes that are present at more than 1% abundance across all samples. **(D) Taxonomic (genus) versus functional (CAZyme) variability across the 37 individuals from 7 host species for each gastrointestinal region**. For each gastrointestinal region, we compute FAVA using relative abundances of either genera (navy, panel A) or CAZYme categories (green, panel C). We compute variability across samples from the 37 host individuals, regardless of host species. Each dot corresponds to the value of FAVA for a gastrointestinal region across the 37 host individuals. The inset panel magnifies the plot for CAZymes.

### Variability in genus abundances

The genus-level compositions of the microbiome samples are shown in Figure 2A. Across host species, all regions of the stomach (2A, blue *x*-axis labels) are dominated by bacteria in the genus *Prevotella*. The samples from the small intestine (2A, yellow *x*-axis labels), on the other hand, are much less homogeneous, with dramatic inconsistency across individuals even within a single region and host species. Samples from the large intestine (2A, red *x*-axis labels) possess a few genera, such as *Bacteroides* (olive), *Clostridium* (navy), and *Ruminococcus* (peach), at similar frequencies across host species and regions of the large intestine.

We first used FAVA to quantify for each region the variability of microbial genus abundances across samples from the same host species (Figure 2B). In order to do this calculation, we first partitioned the 370 samples of microbial genus abundances (37 individuals × 10 regions) into 70 matrices, each corresponding to one of the seven host species and one of the ten gastrointestinal regions. In each matrix, rows represent microbiome samples (vertical bars in Figure 2A) and columns represent microbial genera. We then computed FAVA across the rows of each matrix, quantifying in a single number the variability across all 5 or 6 samples in the same host species and gastrointestinal region.

We find that FAVA is significantly higher in regions of the small intestine than in regions of the other two organs: Wilcoxon rank-sum tests comparing the 21 small-intestine FAVA values (3 small-intestine regions × 7 host species) to the 28 stomach FAVA values (4 stomach regions × 7 host species) or to the 21 large-intestine FAVA values (3 large-intestine regions× 7 host species) have one-sided *P* = 0.002 and *P* < 10^−5^, respectively. FAVA is also lower in large-intestine regions than in stomach regions (Wilcoxon rank-sum test, one-sided *P* = 0.001). These results accord with a view that monitoring microbiome composition via stool sampling alone may obscure substantial among-individual variability present upstream in the digestive tract (Shalon *et al*., 2023; Tropini *et al*., 2017).

Next, we measured the compositional variability of genus abundances for each gastrointestinal region across all host species (Figure 2A, vertical slices delimited by black lines). We partitioned the same 370 samples of microbial genus abundances into 10 matrices, one per gastrointestinal region. Again, matrix rows represent microbiome samples and columns represent microbial genera. We then used FAVA to quantify, for each region, the variability of genus abundances across the 37 individuals from the seven host species (Figure 2D, navy). We observe that compositional variability across the 37 individuals from the 7 host species is highest at the beginning of the small intestine, and lowest in the large intestine, with the stomach possessing an intermediate amount of variability. Variability changes continuously along the gastrointestinal tract: except for a low point at the reticulum, a peak at the duodenum, and a slight dip at the cecum, the FAVA value for each region is between that of its preceding and subsequent regions.

### Variability in microbiome function

The functional profile of a microbial community, measured in terms of the types of genes present, provides information not captured by the community’s taxonomic composition (Louca *et al*., 2018). Abundances of gene functional categories are generated by mapping shotgun metagenomic reads to a database of gene sequences grouped by function. We focus here on 350 carbohydrate-active enzymes (CAZymes), which determine the ability of a microbial community to break down complex carbohydrates (Drula *et al*., 2022) (Figure 2C). In general, we expect to see lower variability in functional categories such as CAZymes than in taxonomic categories such as genera because of the phenomenon of functional redundancy: multiple microbial taxa carry out similar metabolic processes, allowing the taxonomic composition of a community to vary without influencing its function (Turnbaugh *et al*., 2007; Burke *et al*., 2011; Lozupone *et al*., 2012; Oakley *et al*., 2014; Louca *et al*., 2018; Estrela *et al*., 2022). Because FAVA can be computed irrespective of the number of categories, we can use it to compare the variabilities of very different types of data, such as taxonomic and functional abundances.

To quantify functional redundancy in each region of the gastrointestinal tract, we compared the variability of genus abundances across the 37 host individuals to the variability of CAZyme abundances across the same 37 host individuals. We expect functional redundancy to keep CAZyme variability lower than genus variability, with a larger difference between the taxonomic and functional variability indicating stronger functional redundancy. We quantified genus-level taxonomic variability in each of 10 gastrointestinal regions by computing FAVA across vectors of genus abundances sampled from the 37 host individuals, irrespective of host species. This computation resulted in 10 values of FAVA, one per gastrointestinal region (Figure 2D, navy). We repeated this computation with CAZyme abundances in place of genus abundances in order to quantify CAZyme variability across host species in each gastrointestinal region (Figure 2D, green). We used bootstrapping across abundance vectors to compare variability values between pairs of regions, since had only one FAVA value per data type and gastrointestinal region. See the Materials and Methods for details on the bootstrapping procedure.

We see in Figure 2D that values of FAVA for functional data (CAZyme, green) are about one-tenth those of taxonomic data (genus, navy), confirming that functional redundancy in the ruminant microbiome leads to much lower functional than taxonomic variability across host species. We also see in Figure 2D (inset) that, although genus-level variability is much higher in the stomach than the large intestine (pairwise bootstrap comparisons between small-intestine regions and other regions, one-sided *P* < 0.05 for each of 21 pairs), compositional variability in CAZyme abundances is as low in the stomach (blue labels) as in the large intestine (red labels) (pairwise bootstrap comparisons between regions in the stomach and regions in the large intestine, one-sided *P* > 0.1 for each of 12 pairs). This disparity in variability of taxonomic versus functional categories in the stomach could suggest that there is more functional redundancy in the stomach than in the large intestine, in the sense that similar levels of CAZyme variability are obtained from a much greater taxonomic variability in the stomach.

In summary, through our analyses of ruminant microbiomes, FAVA allows us to capture the variability of high-dimensional data in a single number that can be easily compared across regions, species, or data types. The analysis finds that within each host species, both taxonomic and CAZyme community composition are most variable across host individuals in the small intestine.

### Defining weighted FAVA

Our initial definition of FAVA (equation 4) does not account for (1) differential weighting of rows (e.g., weighting based on time or distance between samples) or (2) similarities between columns (e.g., phylogenetic similarity between taxa). We now introduce a weighted version that allows for both uneven weighting of samples and for incorporation of information about the relatedness of taxonomic categories (Figure 3).

**Figure 3:**
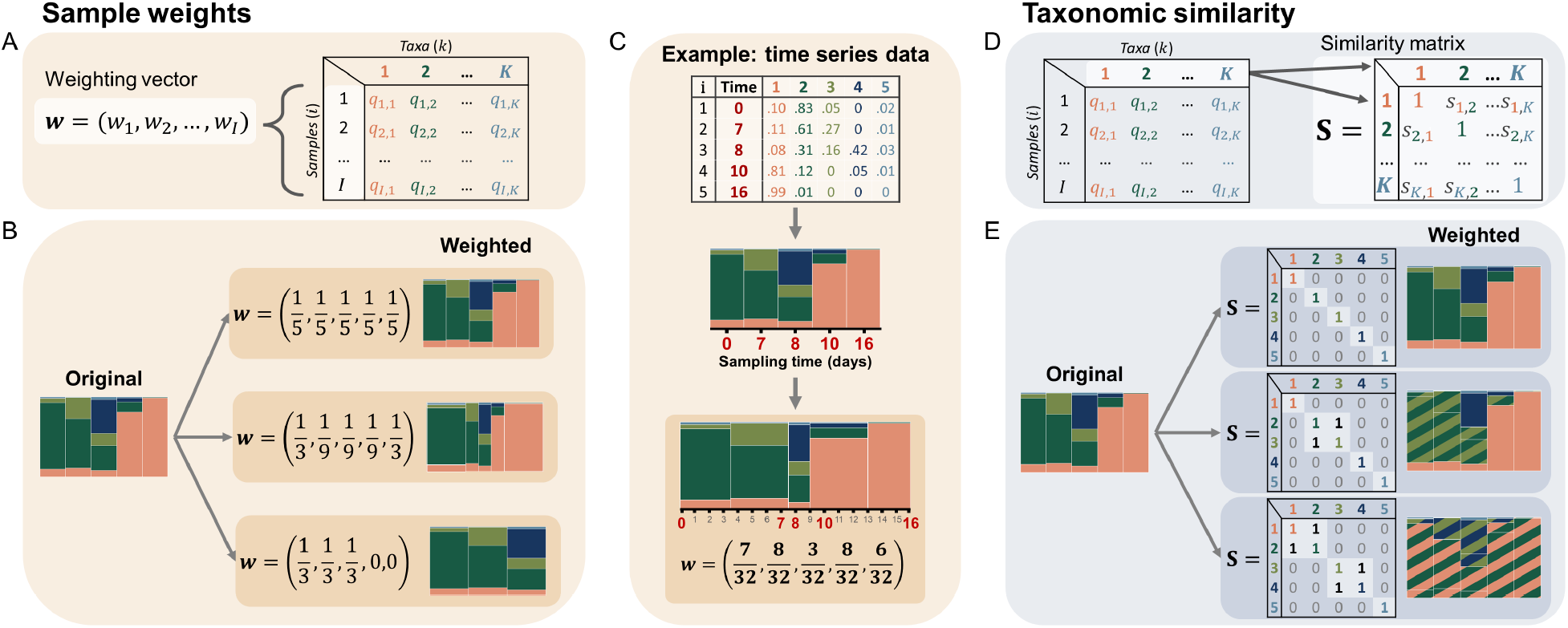
FAVA can incorporate both uneven sample weights and information about the relatedness of taxa. **(A)** FAVA can be weighted by incorporating a normalized weighting vector (**w**). **(B)** Changing the weighting vector, **w**, changes the emphasis placed on each sample by FAVA. The bar plots on the right-hand side represent how FAVA sees the original data under each weighting vector. **(C)** An OTU table with a column representing the collection time of each sample (top) can be visualized as a stacked bar plot, with each bar corresponding to one sample (middle). We can account for uneven sampling times by incorporating a weight vector **w** (bottom) computed using equations 6 and 5. The bottom bar plot represents how FAVA sees these data when this weight vector is used. **(D)** FAVA can also incorporate a similarity matrix (**S**) that represents the relatedness of each pair of taxa. Values can range between 0 and 1, equaling 0 if two taxa are unrelated and 1 if they are identical. **(E)** The coloring of the bar plots on the right-hand side represents how FAVA sees the samples when they are weighted by each similarity matrix. Taxa 2 and 3 are treated as identical in the middle example. In the bottom example, taxa 1 and 2, and taxa 3 and 4 are considered identical. Although we use only zeroes and ones in this schematic, fractional values can be used to represent intermediate levels of similarity.

First, incorporating sample weights is desirable when there is an uneven spatial or temporal distribution of samples, for example if the experimental design includes some weekly samples and some daily samples. In this case, incorporating sample weights allows for greater emphasis on weekly samples, which inform the composition during a seven-day window, than on daily samples (e.g., Figure 3C). Second, incorporating similarity among columns is valuable when the data include some taxa that are closely related and others that are more distant. This weighting helps to make the measure more biologically informed, leading to higher values of FAVA when the taxa that vary in abundance between samples are more distantly related.

We address row weights by incorporating into FAVA a weighting vector **w** = (*w*_1_, *w*_2_, …, *w*_*I*_) that allows for varying emphasis of different samples (Figure 3A-C). Each entry *w*_*i*_ determines the weight placed on sample *i* in the computation of FAVA, and all *w*_*i*_ sum to 1. The default weighting vector assigns identical weight to each sample (Figure 3B, top example). Uneven weights change the emphasis on the different rows; those with larger values contribute more to the diversity calculation (Figure 3B, middle and bottom example). When analyzing time series data, with each sample *i* corresponding to a time *t*_*i*_ between the start, *t*_1_, and the end, *t*_*I*_, a natural choice is to weight each sample *i* by half the distance between the previous sampling time (*t*_*i*−1_) and the subsequent sampling time (*t*_*i*+1_) (equation 5), normalized by the study duration (*T* = *t*_*I*_ − *t*_1_) so that the weights sum to 1 (equation 6). We provide an example of such weights derived from time series data in Figure 3C.

We address similarities among columns by incorporating a similarity matrix, **S** (Figure 3D). For each pair of taxa, this matrix contains a similarity scaled from 0 to 1. Entry *s*_*k,𝓁*_ of the similarity matrix **S** represents the similarity of taxa *k* and *𝓁*: *s*_*k,𝓁*_ = 0 if taxa *k* and *𝓁* are totally dissimilar, *s*_*k,𝓁*_ = 1 if taxa *k* and *𝓁* are identical, and intermediate values represent partial similarity. The diagonal elements of **S** all equal 1, because each taxon is identical to itself. The default similarity matrix is the identity matrix, which has zeroes for all off-diagonal elements (Figure 3E, top example). When ones are placed in off-diagonal elements of the matrix, the corresponding pair of taxa are treated as identical. For example, in the middle example of Figure 3E, taxa 2 and 3 are considered identical, as reflected in the coloring of taxa in the vertical bars to the right. The similarity can be chosen to represent any relevant similarity concept, such as phylogenetic, genetic, or functional similarity.

We explain in the Materials and Methods section how we incorporate both *w*_*i*_ and **S** into equations for Δ_*S*_ and Δ_*T*_ (equations 9 and 10), resulting in an expression for FAVA that considers both uneven row weights and non-trivial column similarities (equation 11). Weighted FAVA (equation 11) reduces to unweighted FAVA (equation 4) when 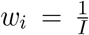 and **S** = **I**_*K*_, a matrix with all *K* diagonal elements equal to 1 and all off-diagonal elements equal to 0 (top examples of Figure 3B and E, respectively).

### Temporal variability and antibiotic perturbation in the human gut microbiome

To demonstrate weighted FAVA as a measure of temporal microbiome variability, we apply it to data from a longitudinal study of gut microbiome composition after antibiotic perturbation (Xue *et al*., 2023). Among 48 subjects, we focused on 22 who took a course of the antibiotic ciprofloxacin midway through the study. For these subjects, stool samples were collected at 26 time points—weekly samples for 9 weeks, as well as daily samples for the three weeks surrounding the antibiotic course (Figure 4A). Xue et al. (Xue *et al*., 2023) inferred the relative abundances of bacterial species over time by shotgun metagenomic sequencing of each sample (Figure 4B, for three of the 22 subjects). We use FAVA to quantify both the impact of the antibiotic perturbation on temporal microbiome variability and the duration of this impact. To account for both the nonuniform sampling timeline and the immense taxonomic diversity of the sampled species, we weight FAVA by both the time intervals between stool samples and the phylogenetic similarity among species. We derive the phylogenetic similarity matrix from an established phylogenetic tree of bacterial species (Nayfach *et al*., 2016), as discussed in the Materials and Methods.

**Figure 4:**
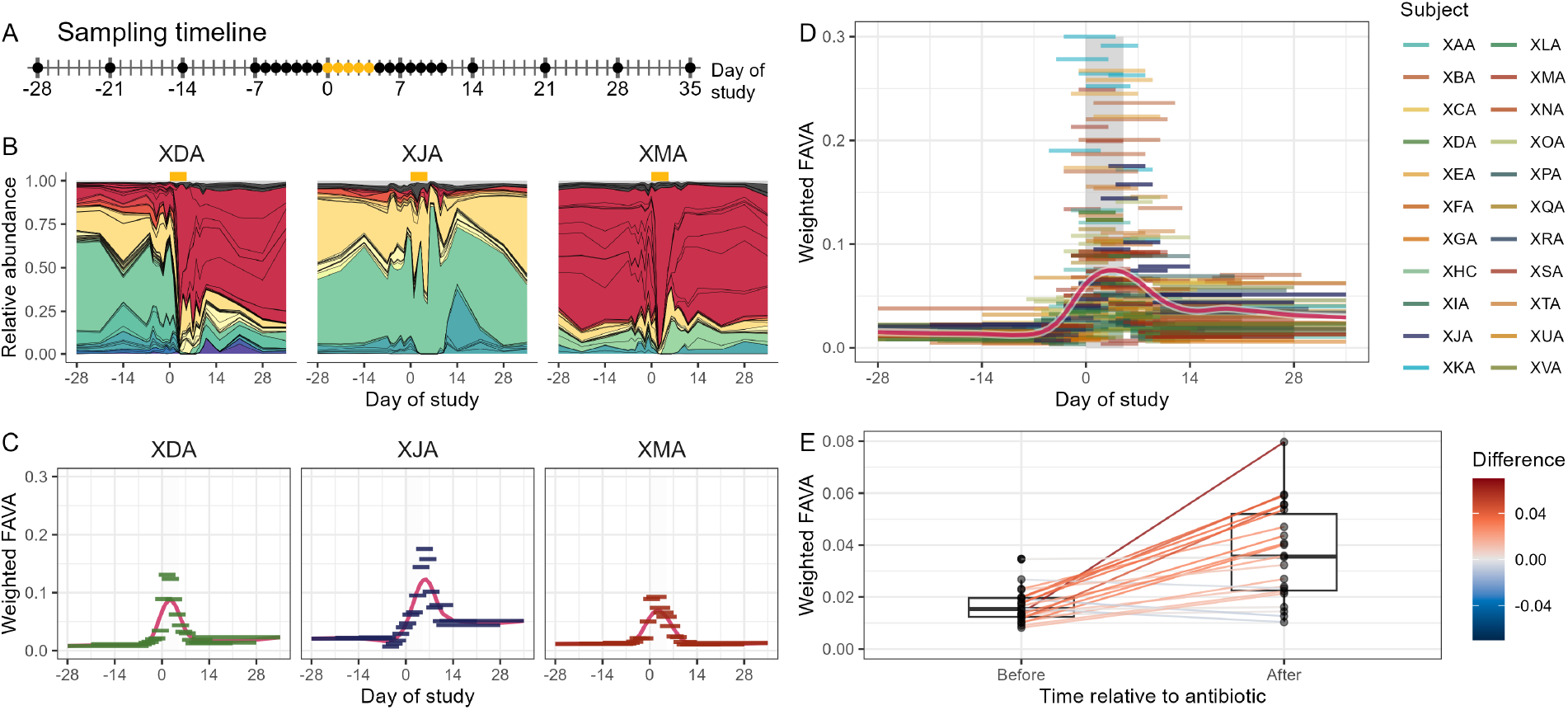
FAVA quantifies changes in the temporal variability of human gut microbiomes following an antibiotic perturbation. **(A) Sampling timeline**. Points correspond to sampling days. Samples are collected weekly for 9 weeks, with daily sampling for three weeks beginning one week before antibiotics. Gold points denote days 0-4, during which the subjects took the antibiotic ciprofloxacin. **(B) Selected relative abundance plots**. Each three-letter code refers to one of 22 subjects. Abundances between sampling times are interpolated by drawing straight lines between abundances for adjacent time points. Colors denote bacterial families and black lines delineate bacterial species. All 22 subjects and the color legend for bacterial families are shown in Figure S1. **(C) Sliding windows of weighted FAVA for selected subjects**. For each subject, we generate sliding windows six samples wide (horizontal bars). The vertical position of each bar is determined by the value of FAVA computed across the six samples in that window. The horizontal breadth of each bar encompasses the time during which the six samples were collected. The vertical gray bar denotes the time during which the subjects were taking antibiotics. The red curve is a smoothing spline fit to all data points with 12 degrees of freedom. Low values of FAVA suggest community composition is stable in a window, whereas high values of FAVA imply temporal variability. **(D) Sliding windows of FAVA for all subjects**. We compute FAVA in six-sample sliding windows for each of the 22 subjects. The red curve is a smoothing spline fit to all data points with 12 degrees of freedom. **(E) Weighted FAVA increases after antibiotic perturbation**. We compute weighted FAVA for each subject either before (days -28 to -1) or after antibiotic perturbation (days 5 to 35), excluding the period during which subjects were taking antibiotics. FAVA is weighted based on both phylogenetic similarity among bacterial species and time between samples. Lines connect values of FAVA for the same subject before and after antibiotic perturbation. Lines are colored according to the difference in FAVA (after minus before). Across all subjects, FAVA increases significantly after antibiotic perturbation (Wilcoxon signed rank test, *P* < 10^−4^).

We first explored how the temporal variability of the gut microbiome changes after an antibiotic perturbation. We find that weighted FAVA is significantly higher after the perturbation than before (Wilcoxon signed-rank test comparing post-antibiotic to pre-antibiotic FAVA values across 22 subjects, *P* < 10^−4^), suggesting that microbiome composition is more temporally variable after antibiotic perturbation (Figure 4E). This result is robust to variability across subjects in the numbers of samples collected in the pre and post-antibiotic time periods (Figure S2).

We explored temporal variability at smaller timescales by computing weighted FAVA in sliding windows throughout the study period. This increased granularity allowed us to quantify changes in temporal variability over the course of the study. In Figure 4D, we show weighted FAVA in sliding windows six samples wide, generating a median of 20 windows per subject. We find that weighted FAVA is significantly higher for the 115 six-sample post-antibiotic windows than for the 89 six-sample pre-antibiotic windows (Wilcoxon rank sum test, *P* < 10^−15^).

The sliding window analysis also allows us to characterize the timeline of the antibiotic perturbation. We find that, for most subjects, the antibiotic perturbation results in a lasting increase in microbiome variability: across the 30 days following completion of the antibiotic, only 8 of the 22 subjects returned to their initial variability level (Figure 4D). However, high levels of variability tend to last for only one or two weeks post-antibiotic: while all subjects began with sub-0.05 FAVA values and 18 of the 22 subjects exceeded 0.05 during the antibiotic period, 11 of these 18 subjects returned to FAVA levels below 0.05 beginning one week post-antibiotic, and 16 of these 18 had sub-0.05 FAVA levels by their final sliding window.

Finally, our sliding window approach allows us to characterize temporal dynamics based on local temporal variability alone. For example, consider subjects XDA, XJA, and XMA, whose variability dynamics are highlighted in Figure 4C. The microbiome of subject XJA does not stabilize postantibiotic, remaining highly variable through the end of the study period. Subjects XDA and XMA, on the other hand, both return to low variability levels within seven days of the conclusion of antibiotics. However, Figure 4B reveals that these subjects represent two different responses to the antibiotic perturbation. Whereas subject XMA returned to the original compositional state after the antibiotic perturbation, subject XDA settled at a new compositional state very different from the initial microbial community.

This example highlights that, by computing FAVA on small windows, we can identify periods of temporal stability in microbiome composition, even when the microbiome has stabilized at a compositional state different from its initial state. For this reason, FAVA complements pairwise distance measures that capture compositional dissimilarity but not temporal variability, such as the Jensen-Shannon divergence, Bray-Curtis dissimilarity, and Jaccard index.

### R Package

We have implemented FAVA in an R package, titled *FAVA*, which is available at github.com/MaikeMorrison/FAVA. The *FAVA* R package contains a function that can compute FAVA, weighted FAVA, and FAVA normalized by the upper bound given the abundance of the most abundant taxon. It also has functions to compute these three versions of FAVA in sliding windows and to visualize the sliding window results in plots such as those in Figure 4C and D. The FAVA package can also visualize relative abundance data in stacked bar plots, and it can statistically compare groups of samples with bootstrapping.

## Discussion

We have introduced an index to quantify variability across many samples of microbiome composition. We defined the measure through an analogy with the population-genetic statistic *F*_*ST*_, considering microbiome samples in place of populations and microbial taxa in place of alleles. FAVA, an *F*_*ST*_ - based Assessment of Variability across vectors of relative Abundances, equals 0 if and only if all microbiome samples are identical, and equals 1 if and only if each microbiome sample contains only a single taxon and there is more than one taxon present across all samples (Figure 1). FAVA can be used as a measure of compositional variability across time points, spatial sampling locations, host individuals, or replicates, quantifying the temporal variability, spatial heterogeneity, or replicability of microbial communities. Because FAVA takes values between 0 and 1 regardless of the number of sampled taxa, we can compare values of FAVA between very different data sets, such as data on abundances of different taxonomic categories.

To demonstrate FAVA’s performance as a measure of microbiome variability across many samples, we analyzed two microbiome data sets: an investigation of ruminant microbiome composition along the gastrointestinal tract (Xie *et al*., 2021), and a longitudinal study of human gut microbiome composition before and after an antibiotic perturbation (Xue *et al*., 2023). In the ruminant data, we found that compositional variability across individuals—either within a host species or across host species—was consistently lower at the end of the gastrointestinal tract than in the middle, supporting the view that substantial inter-individual heterogeneity is missed when microbiomes are monitored by fecal sampling alone (Figure 2B and D) (Shalon *et al*., 2023; Tropini *et al*., 2017). We found that, in all gastrointestinal regions, taxonomic abundances were much more variable across individuals than were functional abundances, a result that corroborates observations of microbial functional redundancy in the gastrointestinal tract (Figure 2D) (Louca *et al*., 2018).

In the human microbiome data, we found that antibiotic perturbations significantly destabilize microbial communities, resulting in elevated temporal variability following an antibiotic (Figure 4E). Computing FAVA in sliding windows across temporal samples for each subject increased the granularity of this analysis. We found that, although elevated variability lasted for only 1-2 weeks post-antibiotic on average, few subjects returned to their pre-antibiotic variability levels during the study duration (Figure 4C and D). We also highlighted FAVA’s ability to quantify temporal variability separate from compositional state by focusing on subjects XDA and XMA, who returned to their pre-antibiotic variability levels (Figure 4C) even though only subject XMA returned to the original composition (Figure S3).

We introduced two extensions of FAVA: weighted FAVA (equation 11), which can incorporate both similarity among taxa and distance between samples into the computation, and normalized FAVA, which accounts for the abundance of the most abundant taxon, allowing for more meaningful measurement of variability across small numbers of samples. In our analysis of human gut microbiome data over time (Xue *et al*., 2023), the use of weighted FAVA helped to account for both the combination of weekly and daily samples and the broad range of species appearing in the data.

FAVA values can be influenced by the choice of weights. For example, Figure S4 presents two simple hypothetical OTU tables with a difference in FAVA of nearly 0.5 when weighted by taxonomic similarity, despite having identical unweighted FAVA values. Nevertheless, in our analysis of human microbiome data, although individual FAVA values shift with the incorporation of weights, FAVA values computed across post-antibiotic samples are consistently higher than those computed across pre-antibiotic samples, regardless of whether FAVA is weighted by sampling times, taxonomic similarity, or both (Figure S5).

Our measure, which we have implemented in an R package, contributes to a large body of methods for the analysis of microbiome relative abundance data (McMurdie and Holmes, 2013; Bolyen *et al*., 2019). We emphasize, however, that FAVA is a multi-sample compositional variability measure, setting it apart from the many existing measures of pairwise compositional similarity, such as Unifrac, Bray-Curtis dissimilarity, and the Jensen-Shannon divergence (Figure 1A) (Bray and Curtis, 1957; Lin, 1991; Lozupone and Knight, 2005). For example, two separate collections of microbiome samples can have identical values of FAVA, but wildly different mean compositions (e.g., Figure 4B and C). Similar values of FAVA therefore reflect similarities in the spatial or temporal dynamics shaping variability, not compositional similarity. FAVA builds on a rich literature of population-genetic and ecological frameworks for hierarchical partitioning of genetic, taxonomic, and phylogenetic diversity across individuals and communities (Lewontin, 1972; Lande, 1996; Ricotta, 2005; Hardy and Senterre, 2007; Ellison, 2010). Indeed, *F*_*ST*_ has sometimes been used as a measure of compositional variability in ecological contexts (Gilbert and Levine, 2017). Future applications of FAVA can span the range of questions that researchers pose about compositional variability, from understanding temporal variability in infant microbiomes (Koenig *et al*., 2011; Yassour *et al*., 2016) to quantifying the repeatability of community assembly across experimental replicates to identifying the timing of compositional stability in serial passaging experiments (Goldford *et al*., 2018; Estrela *et al*., 2022). Because FAVA measures a fundamentally different phenomenon relative to existing methods for microbiome analysis, it opens up a suite of novel research questions relating to temporal stability, individual heterogeneity, spatial variability, and replicability.

## Materials and Methods

### Notation

*Q* denotes an OTU table with *I* rows, each representing a microbiome sample, and *K* columns, each representing a microbial species, OTU, genus, functional unit, or other such category. Entry *q*_*i,k*_ represents the relative abundance of taxon *k* in sample *i*. Each row must sum to 1. We use “sample *i*” to refer to the *i*^*th*^ row of *Q* (*q*_*i*,*_).

### Bootstrapping protocol

We use bootstrapping (Efron and Tibshirani, 1993) to determine whether two values of FAVA are significantly different. Consider two OTU tables, *A* with *n* rows and *B* with *m* rows. The observed difference in FAVA values between these two matrices is *D*_obs_ = *F*_*ST*_ (*A*) − *F*_*ST*_ (*B*). Our null hypothesis is that there is no difference in *F*_*ST*_ values between the communities sampled to form tables *A* and *B*.

To test this hypothesis, we first merge the two OTU tables into a single matrix, *Q*_null_, which has *n* + *m* samples corresponding to the samples in *A* and *B*. We then randomly draw *n* or *m* rows with replacement from *Q*_null_ to generate bootstrap replicates for the *A* and *B, A*_boot_ and *B*_boot_ respectively. Finally, we compute the difference in FAVA values between these bootstrap replicate matrices, *D*_boot_ = *F*_*ST*_ (*A*_boot_) − *F*_*ST*_ (*B*_boot_). Repeating this procedure many times (e.g., 1,000 times) to generate many values of *D*_boot_ results in a bootstrap distribution of differences in FAVA values between *A* and *B*.

We test our null hypothesis that there is no difference in FAVA values between *A* and *B* by comparing the observed difference, *D*_obs_, to the bootstrap distribution of differences. We obtain a one-sided P-value by computing the proportion of bootstrapped differences *D*_boot_ that are either greater than or less than the observed difference *D*_obs_. We obtain a two-sided P-value by comparing |*D*_boot_| to |*D*_obs_|. A worked example of this computation is available in the *FAVA* R package vignette.

### Defining weighted *F*_*ST*_

#### Incorporating uneven sample weights

For each sample *i* = 1, …, *I*, we choose a weight *w*_*i*_ ⩾ 0 such that 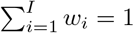. To evenly weight all samples, choose 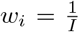 for all *i*. Uneven weights *w*_*i*_ can be chosen to account for properties such as sample size or the spatial or temporal distance between samples. If samples come from a time series, with *t*_*i*_ representing the sampling time of sample *i*, we recommend defining 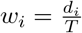 (equation 6), where *T* = *t*_*I*_ − *t*_1_ is the study duration and *d*_*i*_ is half the time from the sample before *i* to the sample after *i* (equation 5):

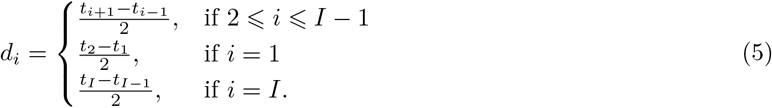

Because 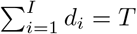

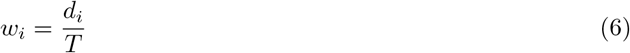

is a weight that sums to 1 over all *i* and represents the proportion of the study duration accounted for by sample *i*. Note that in the case of evenly spaced time samples, under the weighting 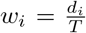, the first and last sample are given half as much weight as the intermediate samples. This means that the uniform case is similar to but not exactly equal to the original, unweighted definition of *F*_*ST*_, which has 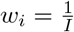.

A standard definition for *F*_*ST*_ is *F*_*ST*_ = (Δ_*T*_ − Δ_*S*_)*/*Δ_*T*_ (equation 4), where Δ_*S*_ is the mean sample Gini-Simpson diversity and Δ_*T*_ is the total Gini-Simpson diversity (equations 2, 3):

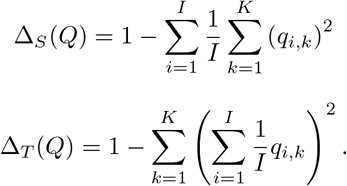

We incorporate time information by replacing the uniform weights 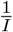 with not necessarily uniform weights **w** = (*w*_1_, *w*_2_, …, *w*_*I*_):

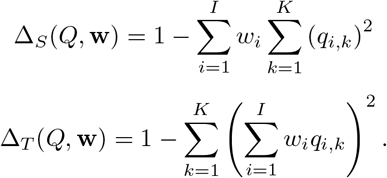

*F*_*ST*_ (*Q*, **w**) = (Δ_*T*_ (*Q*, **w**) − Δ_*S*_(*Q*, **w**))*/*Δ_*T*_ (*Q*, **w**) is thus a definition of *F*_*ST*_ that allows for uneven weighting of samples. Note that this weighting can account for differences in spacing between samples, but not for differences in relative ordering of samples.

#### Incorporating taxonomic similarity

In addition to incorporating uneven sample weights, we may wish to account for the similarity between taxa. We capture information about the similarity among all *K* taxa through the symmetric, *K* × *K* similarity matrix **S**. The entry in row *k* and column *𝓁* of **S**, *s*_*k,𝓁*_, represents the similarity between taxon *k* and taxon *𝓁*. Diagonal elements satisfy *s*_*k,k*_ = 1 because each taxon is identical to itself, and we define the similarity between identical taxa to be 1. Off-diagonal elements take values in [0, 1], equaling 0 if two taxa are minimally similar, and 1 if they are identical. If **S** is the identity matrix (i.e., *s*_*k,𝓁*_ = 0 for all *k ≠ 𝓁*), then all distinct taxa are treated as minimally similar and our weighted version of *F*_*ST*_ must reduce to its original, unweighted definition (equation 4).

In order to incorporate **S** into the definition of *F*_*ST*_, we first introduce Leinster and Cobbold’s (Leinster and Cobbold, 2012) idea of “mean ordinariness” across taxa in a microbiome sample. The “ordinariness” of taxon *k* in sample *i* is the mean similarity between that taxon and every other taxon in the sample, weighted by the taxon abundances. It is computed for each taxon by multiplying the similarity matrix (**S**) by sample *i* (*q*_*i*,*_). This computation produces a vector whose *k*^*th*^ entry, 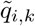_*i,k*_, represents the mean similarity between species *k* and a random taxon from sample *i*:

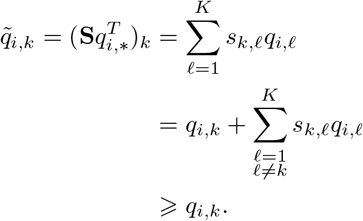

In other words, 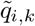 measures the ordinariness of taxon *k* within sample *i*. On one extreme,

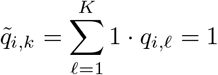

if taxon *k* is identical to all other taxa in sample *i*. In this case, taxon *k* is maximally ordinary in relation to the other taxa in the sample. On the other extreme,

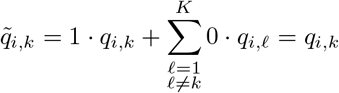

if taxon *k* has similarity 0 to all other taxa in sample *i*. In this case, taxon *k* is minimally ordinary in relation to the other sampled taxa. The mean taxon ordinariness across all taxa in sample *i*, weighted by their abundances, is

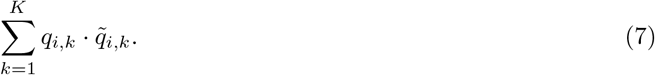

This quantity has been explored in previous work on ecological diversity indices (Leinster and Cobbold, 2012). It is large (i.e., approaching or equal to 1) if the sample is concentrated in a few very similar taxa, whereas it is small (i.e., approaching 0) if the sample is spread across many unrelated taxa. If **S** is the identity matrix, with ones along the diagonal and zeroes for off-diagonal elements, then 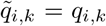 for all *k* and equation 7 reduces to the mean taxon abundance across all taxa in sample 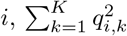.

We proceed by extending this idea of mean ordinariness into the framework of *F*_*ST*_. First, recall the original definition of the Gini-Simpson index (equation 1), which can be interpreted as one minus the mean taxon abundance across all taxa in a sample:

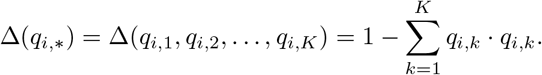

We incorporate the similarity matrix **S** into the Gini-Simpson index by replacing the mean abundance across taxa, 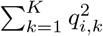, with the mean ordinariness across taxa, 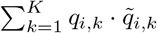, giving the following definition:

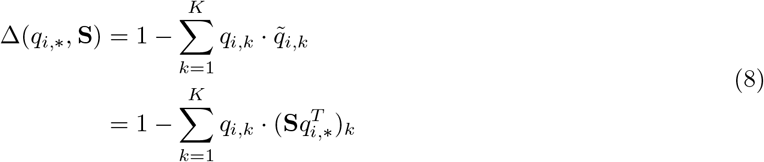

Equation 8 reduces to equation 1 if **S** is the identity matrix. In this case, each taxon is considered extraordinary, with similarity 0 to all other taxa. However, if **S** has non-zero off-diagonal elements, equation 8 is able to account for the similarity among taxa in its computation of diversity.

Finally, we extend equation 8 to define versions of Δ_*S*_ and Δ_*T*_ :

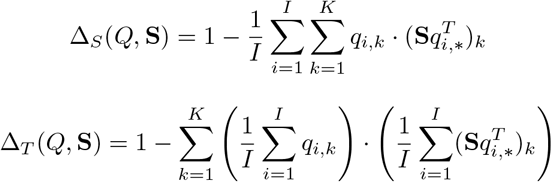

Using these extensions of Δ_*S*_ and Δ_*T*_ in a computation of *F*_*ST*_ yields a compositional variability measure that accounts for taxonomic similarity: *F*_*ST*_ (*Q*, **S**) = (Δ_*T*_ (*Q*, **S**) − Δ_*S*_(*Q*, **S**))*/*Δ_*T*_ (*Q*, **S**).

#### Simultaneously incorporating uneven row weights and taxonomic similarity

We simultaneously incorporate both **S** and **w** into equations for Δ_*S*_ and Δ_*T*_ in order to develop a compositional variability statistic that accounts for both weighting of samples and similarity among taxa (equations 9, 10, 11):

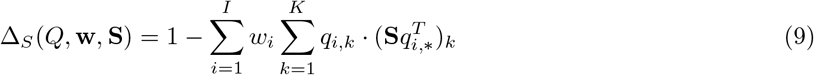

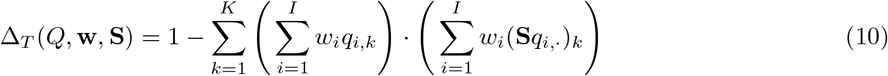

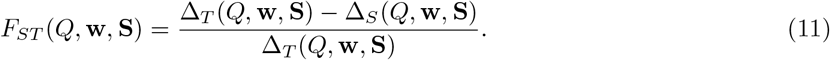

If *w*_*i*_ = 1*/I* for all *i*, and **S** is the identity matrix, this weighted definition of *F*_*ST*_ (equation 11) reduces to the unweighted version of *F*_*ST*_ (equation 4).

#### Computing the phylogenetic similarity matrix

In our analysis of human microbiome data (Figure 4), we chose to weight FAVA by the phylogenetic similarity among the sampled bacterial species. We computed the phylogenetic similarity matrix through a two-step process. First, we computed the patristic distance between each pair of sampled bacterial species based on a microbial phylogeny from Nayfach et al. (Nayfach *et al*., 2016). We performed this computation with the “cophenetic.phylo” function in the *ape* R package (Paradis and Schliep, 2019). Second, we transformed the pairwise patristic distances, which range from 0 for identical species to ∼3.7 for very distantly related species, to similarities, which range from 0 for very distantly related species to 1 for identical species. We chose to convert the patristic distance between species *k* and *𝓁* (*d*_*k,𝓁*_) to a similarity (*s*_*k,𝓁*_) using the exponential transformation *s*_*k,𝓁*_ = exp (−*d*_*k,𝓁*_).

Different transformations of distances to similarities result in different distributions of similarity values. In our case, the similarity matrix computed with the exponential transformation had a median value of 0.087, with first and third quartiles [0.067, 0.119]. Other transformations are defensible as well. The linear difference transformation (*s*_*k,𝓁*_ = 1 − *d*_*k,𝓁*_*/* max *d*_*k,𝓁*_), for example, yields a mean value of 0.716, with with first and third quartiles [0.685, 0.752]. We note that the main results of Figure 4 do not depend on the choice of transformation. In particular, regardless of the transformation used, FAVA values increase during the antibiotic perturbation and are significantly higher post-antibiotic than pre-antibiotic (Figure S6).

### Datasets

#### Ruminant data

In our first data example, we analyzed genus and CAZyme abundances inferred from metagenomic sequencing of samples collected at 10 gastrointestinal regions from 37 ruminant host individuals representing 7 host species. This data set was collected and published by Xie et al. (Xie *et al*., 2021). We downloaded the data from: http://rummeta.njau.edu.cn/rumment/resource/metagenomicsPage. The genus abundances were found in the file “RGMGC.genus.profile.txt” which was available for download under the heading “Genus profile (genus abundance profile table for 370 GIT samples).” The CAZyme abundances were found in the file “RGMGC.cazy.profile.family.txt,” which was available for download under the heading “Cazy profile (Cazy abundance profile table for 370 GIT samples).” For both genera and CAZymes, the published data contained absolute abundances. We converted absolute abundances to relative abundances before performing our analyses.

### Human microbiome data

In our second data example, we analyzed data generated by Xue et al. (Xue *et al*., 2023).

## Acknowledgements

M.L.M. acknowledges support from an NSF graduate research fellowship and a Stanford graduate fellowship. K.S.X. acknowledges support from a James McDonnell Foundation Postdoctoral Fellowship in Understanding Dynamic and Multi-Scale Systems, a Jane Coffin Childs Memorial Fund Postdoctoral Fellowship, and NIH/NIAID grant R21-AI168860. N.A.R. acknowledges support from NIH grant R01 HG005855. We thank Nicolas Alcala, Jessica Grembi, Po-Yi Ho, Egor Lappo, and Chloe Shiff for helpful comments.

## Supplemental Materials

**Figure S1:**
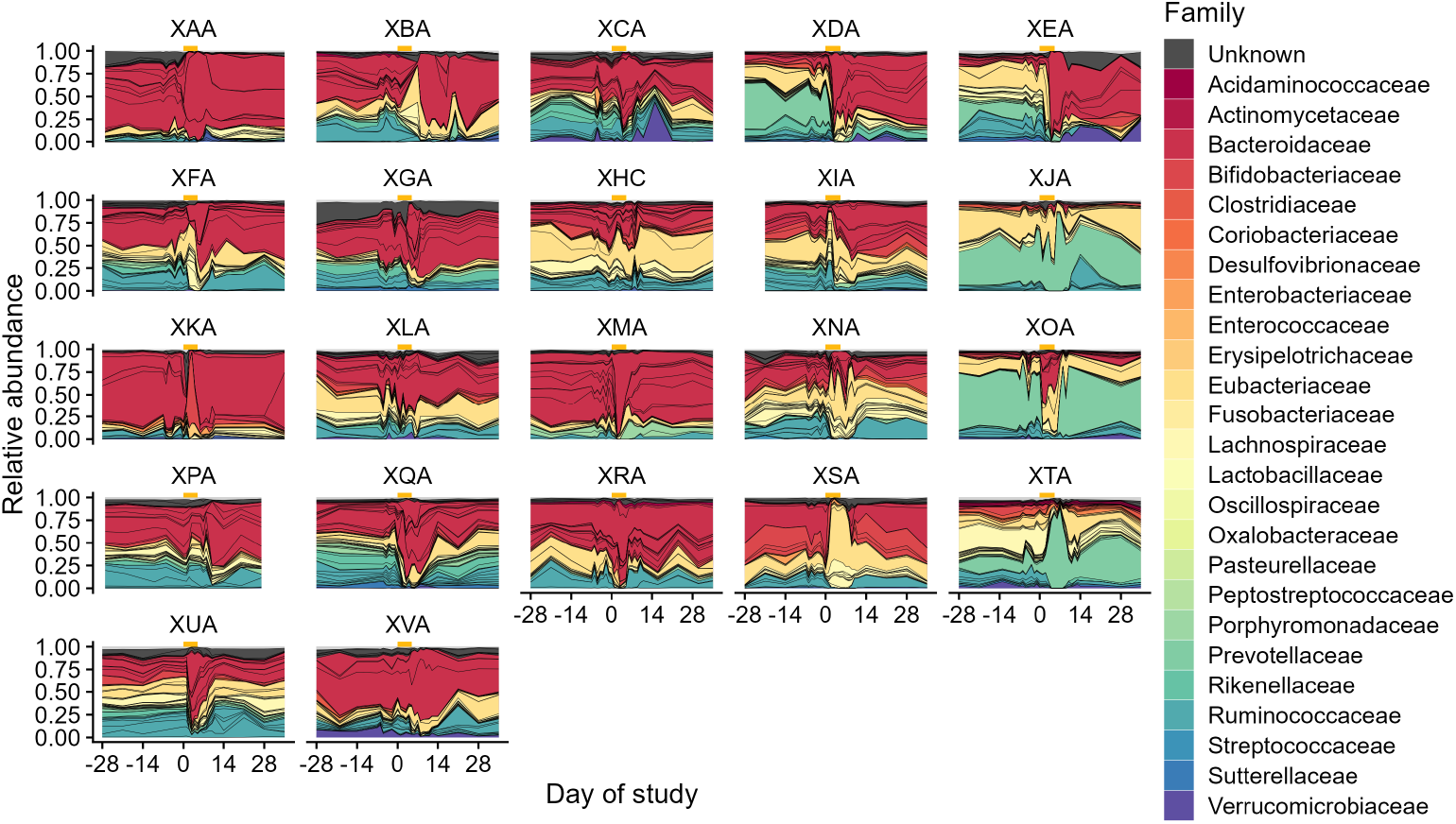
Relative abundances of bacterial genera over time for 22 subjects. Each three-letter code refers to one subject. Gold bars denote days 0-4, during which the subjects took the antibiotic ciprofloxacin. Samples are collected once weekly for 9 weeks, with daily sampling for three weeks beginning one week before antibiotics. Abundances between sampling times are interpolated by drawing straight lines between abundances for adjacent time points. Colors denote bacterial families and black lines delineate bacterial species. Species are identified by mapping metagenomic sequencing reads to a database of single-copy bacterial genes. The figure is adapted from Xue et al. Xue *et al*. (2023).

**Figure S2:**
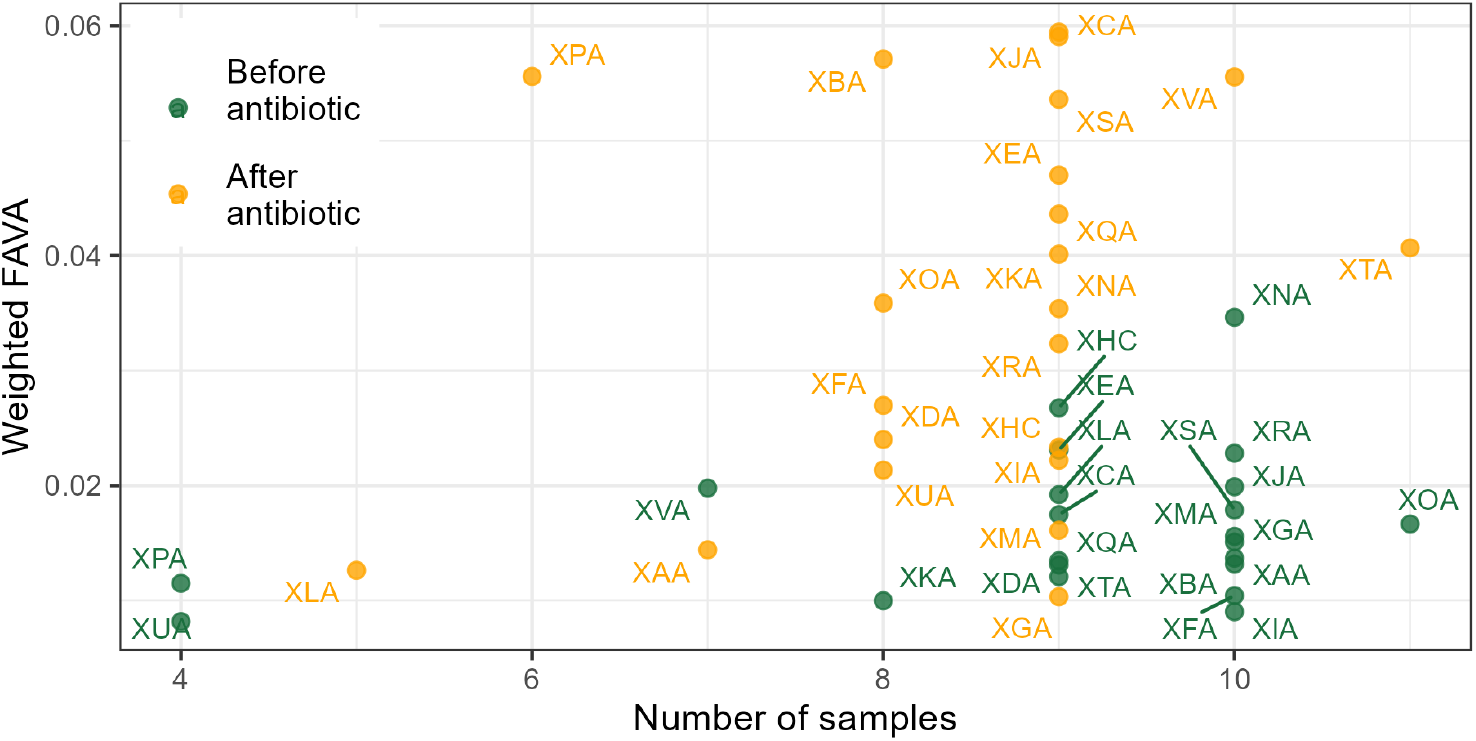
Change in weighted FAVA before and after the antibiotic perturbation is driven by antibiotic, not by the number of samples before or after antibiotic. Each point represents an analysis of all samples collected from one of 22 subjects either before the antibiotic perturbation (green points; samples taken between day -28 and day -1) or after the antibiotic perturbation (orange points; samples taken between day 5 and day 35). The *x*-axis gives the total number of samples analyzed for each point, and the *y*-axis gives the value of FAVA computed across these samples. The y-axis presents the same data shown in Figure 4E. While subjects tended to have more samples before than after the antibiotic perturbation (Wilcoxon signed rank test, one-sided *P* = 0.013), there was no correlation between FAVA and the number of samples included in the computation (Spearman correlation *ρ* = −0.001, *P* = 0.992). We therefore conclude that the difference in FAVA values observed in Figure 4E is driven by the antibiotic perturbation, not by differences in the numbers of samples included in computations of FAVA.

**Figure S3:**
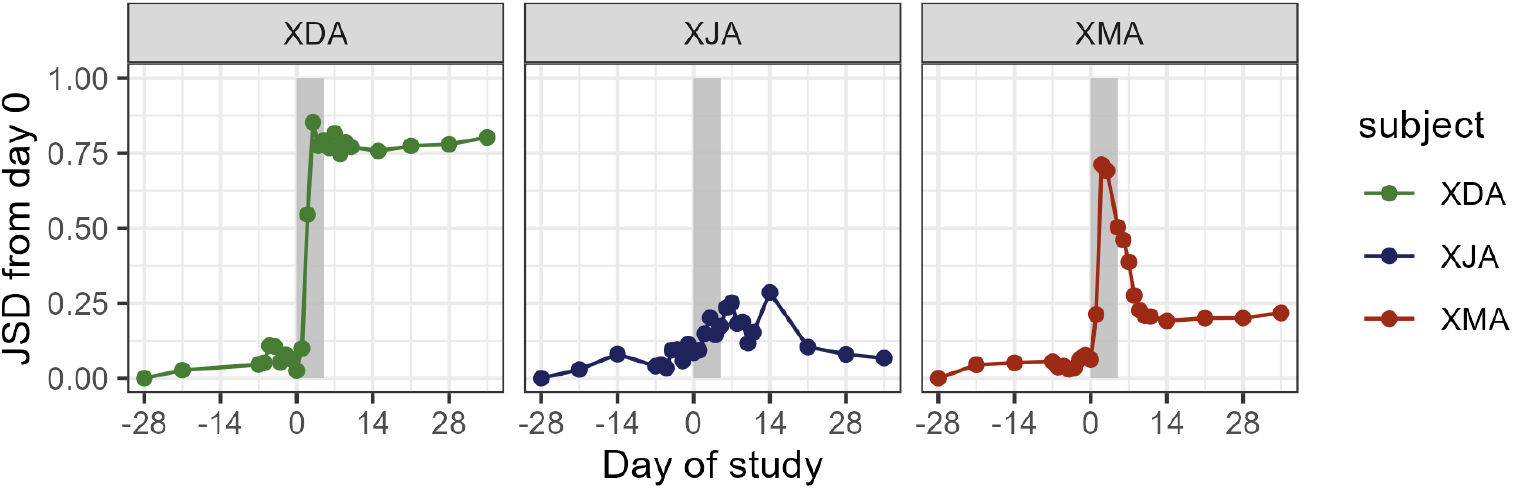
The Jensen-Shannon divergence (JSD) from the initial timepoint captures compositional changes in microbial communities following antibiotic perturbation (grey bar). JSD is computed using species relative abundances.

**Figure S4:**
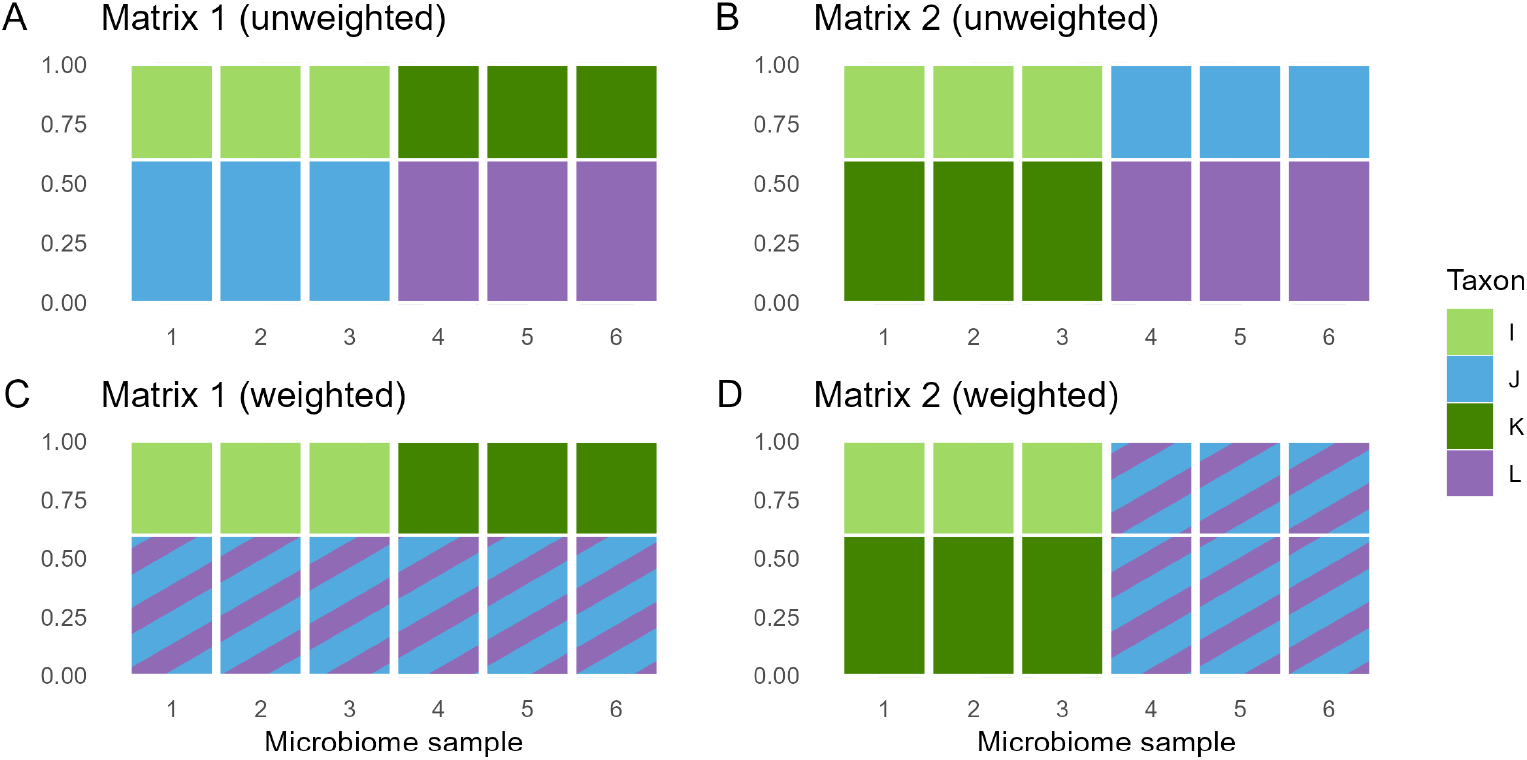
The effect of incorporating taxonomic similarity into the FAVA computation depends on the taxonomic composition of the sampled communities. Each panel shows one matrix representing the compositions of six hypothetical microbiome samples. Without accounting for taxonomic similarity, both Matrix 1 (A) and Matrix 2 (B) have FAVA values of 0.35. We consider a similarity matrix that treats taxon J and taxon L as identical (similarity equal to 1), and all other distinct taxa as totally unrelated (similarity equal to 0). Incorporating this similarity matrix in the computation of FAVA decreases Matrix 1’s FAVA value to 0.14 (C) and increases Matrix 2’s FAVA value to 0.61 (D).

**Figure S5:**
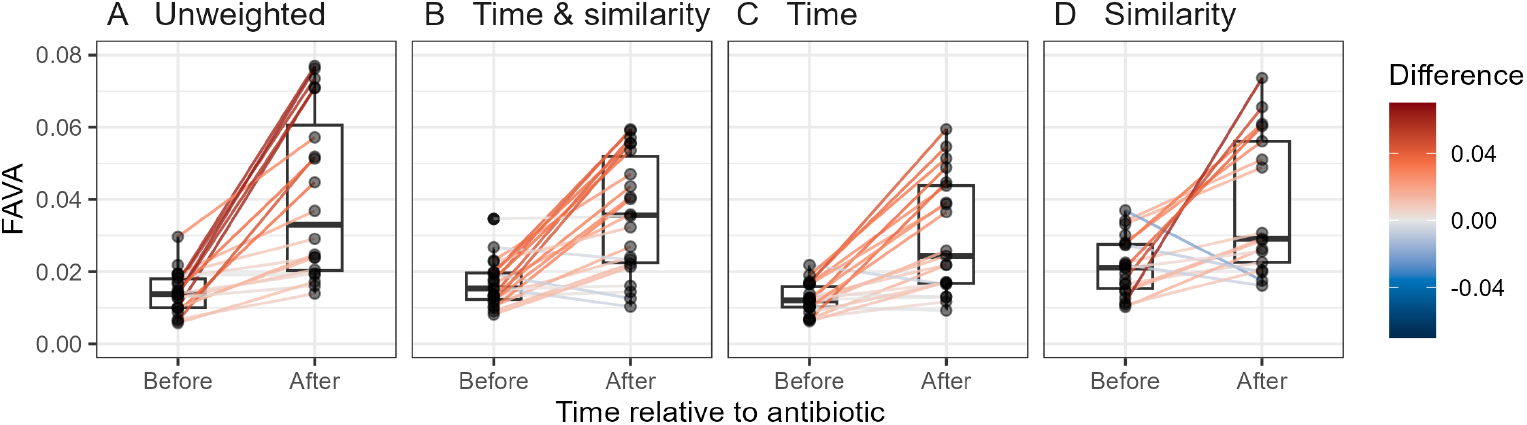
FAVA increases after antibiotic perturbation regardless of FAVA weighting. We compute FAVA for each subject either before the antibiotic perturbation (samples in days -28 to -1) or after the antibiotic perturbation (samples in days 5 to 35). Lines connect values of FAVA for the same subject before and after the antibiotic. The four panels correspond to four different types of weighting for the FAVA measure: **(A)** No weights. **(B)** Weighting by both time between samples and phylogenetic similarity among taxa. **(C)** Weighting just by time. **(D)** Weighting just by phylogenetic similarity. In all four panels, FAVA is significantly higher after the antibiotic perturbation than before (Wilcoxon signed rank test, *P* < 10^−4^ for each panel).

**Figure S6:**
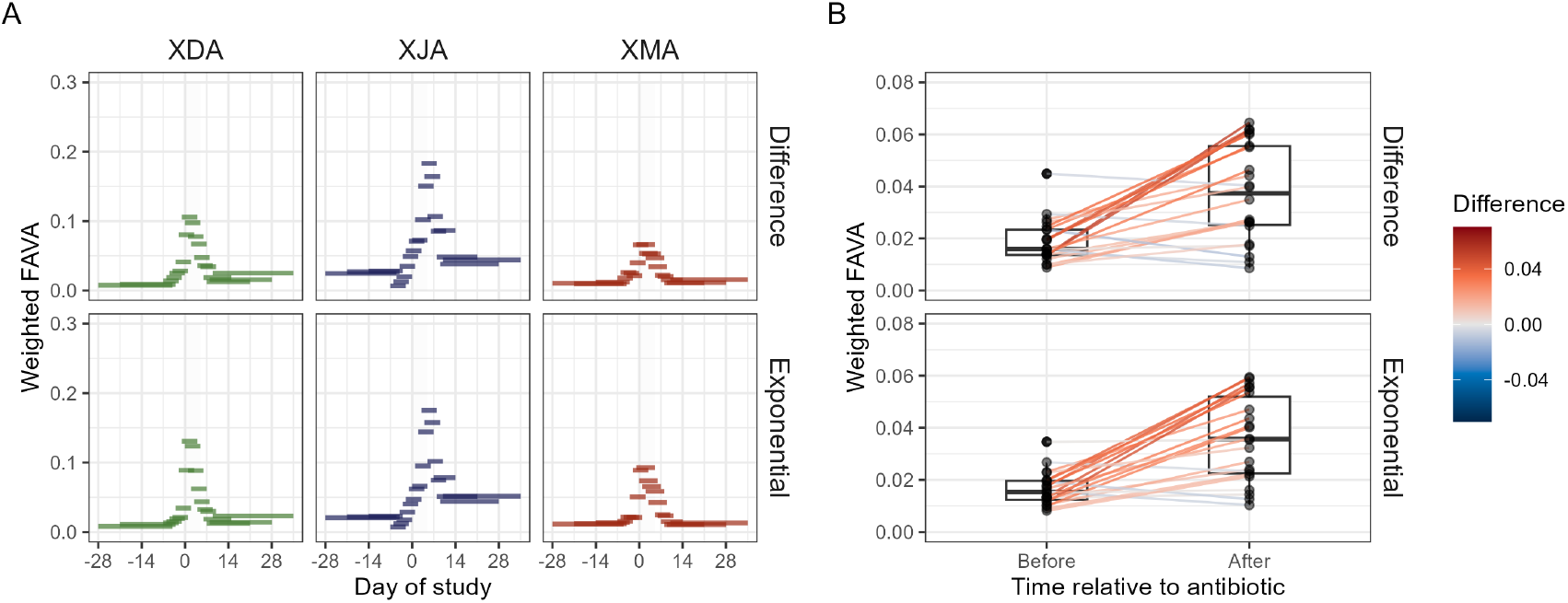
Comparing results from Figure 4 when using different similarity matrices. Panel A reproduces Figure 4C; panel B reproduces Figure 4E. The top row of each panel gives the results using a similarity matrix defined from the patristic distance matrix using the linear difference transformation. The bottom row of each panel gives the results using the exponential transformation, which are the results presented in the main text. The results are largely consistent between the two transformations. For example, as is true for the exponential results presented in the main text, FAVA values computed with the similarity matrix generated using the linear difference transformation are higher after antibiotic perturbation than before (Wilcoxon signed-rank test comparing post-antibiotic to pre-antibiotic FAVA values across subjects, *P* = 0.0003).

## References

Alcala, N., and N. A. Rosenberg, 2022 Mathematical constraints on FST: Multiallelic markers in arbitrarily many populations. Philosophical Transactions of the Royal Society B: Biological Sciences 377: 20200414.

Bashan, A., T. E. Gibson, J. Friedman, V. J. Carey, S. T. Weiss, et al., 2016 Universality of human microbial dynamics. Nature 534: 259–262.

Bolyen, E., J. R. Rideout, M. R. Dillon, N. A. Bokulich, C. C. Abnet, et al., 2019 Reproducible, interactive, scalable and extensible microbiome data science using QIIME 2. Nature Biotechnology 37: 852–857.

Bray, J. R., and J. T. Curtis, 1957 An ordination of the upland forest communities of southern Wisconsin. Ecological Monographs 27: 325–349.

Burke, C., P. Steinberg, D. Rusch, S. Kjelleberg, and T. Thomas, 2011 Bacterial community assembly based on functional genes rather than species. Proceedings of the National Academy of Sciences of the United States of America 108: 14288–14293.

Coyte, K. Z., J. Schluter, and K. R. Foster, 2015 The ecology of the microbiome: Networks, competition, and stability. Science 350: 663–666.

David, L. A., A. C. Materna, J. Friedman, M. I. Campos-Baptista, M. C. Blackburn, et al., 2014 Host lifestyle affects human microbiota on daily timescales. Genome Biology 15: 1–15.

Dethlefsen, L., and D. A. Relman, 2011 Incomplete recovery and individualized responses of the human distal gut microbiota to repeated antibiotic perturbation. Proceedings of the National Academy of Sciences of the United States of America 108: 4554–4561.

Drula, E., M. L. Garron, S. Dogan, V. Lombard, B. Henrissat, et al., 2022 The carbohydrate-active enzyme database: Functions and literature. Nucleic Acids Research 50: D571–D577.

Efron, B., and R. Tibshirani, 1993 An Introduction to the Bootstrap. Chapman and Hall, New York.

Ellison, A. M., 2010 Partitioning diversity. Ecology 91: 1962–1963.

Estrela, S., J. C. Vila, N. Lu, D. Bajić, M. Rebolleda-Gómez, et al., 2022 Functional attractors in microbial community assembly. Cell Systems 13: 29–42.e7.

Faith, J. J., J. L. Guruge, M. Charbonneau, S. Subramanian, H. Seedorf, et al., 2013 The long-term stability of the human gut microbiota. Science 341: 1237439.

Fassarella, M., E. E. Blaak, J. Penders, A. Nauta, H. Smidt, et al., 2021 Gut microbiome stability and resilience: Elucidating the response to perturbations in order to modulate gut health. Gut 70: 595–605.

Flores, G. E., J. G. Caporaso, J. B. Henley, J. R. Rideout, D. Domogala, et al., 2014 Temporal variability is a personalized feature of the human microbiome. Genome biology 15: 531.

Folz, J., R. N. Culver, J. M. Morales, J. Grembi, G. Triadafilopoulos, et al., 2023 Human metabolome variation along the upper intestinal tract. Nature Metabolism 5: 777–788.

Gilbert, B., and J. M. Levine, 2017 Ecological drift and the distribution of species diversity. Proceedings of the Royal Society B: Biological Sciences 284: 20170507.

Goldford, J. E., N. Lu, D. Bajić, S. Estrela, M. Tikhonov, et al., 2018 Emergent simplicity in microbial community assembly. Science 361: 469–474.

Guthrie, L., S. P. Spencer, D. Perelman, W. Van Treuren, S. Han, et al., 2022 Impact of a 7-day homogeneous diet on interpersonal variation in human gut microbiomes and metabolomes. Cell Host and Microbe 30: 863–874.e4.

Hardy, O. J., and B. Senterre, 2007 Characterizing the phylogenetic structure of communities by an additive partitioning of phylogenetic diversity. Journal of Ecology 95: 493–506.

Ji, B. W., R. U. Sheth, P. D. Dixit, Y. Huang, A. Kaufman, et al., 2019 Quantifying spatiotemporal variability and noise in absolute microbiota abundances using replicate sampling. Nature Methods 16: 731–736.

Jost, L., P. Devries, T. Walla, H. Greeney, A. Chao, et al., 2010 Partitioning diversity for conservation analyses. Diversity and Distributions 16: 65–76.

Kang, J. T., J. J. Teo, D. Bertrand, A. Ng, A. Ravikrishnan, et al., 2022 Long-term ecological and evolutionary dynamics in the gut microbiomes of carbapenemase-producing Enterobacteriaceae colonized subjects. Nature Microbiology 7: 1516–1524.

Koenig, J. E., A. Spor, N. Scalfone, A. D. Fricker, J. Stombaugh, et al., 2011 Succession of microbial consortia in the developing infant gut microbiome. Proceedings of the National Academy of Sciences of the United States of America 108: 4578–4585.

Kurokawa, K., T. Itoh, T. Kuwahara, K. Oshima, H. Toh, et al., 2007 Comparative metagenomics revealed commonly enriched gene sets in human gut microbiomes. DNA Research 14: 169–181.

Lande, R., 1996 Statistics and partitioning of species diversity, and similarity among multiple communities. Oikos 76: 5.

Leinster, T., and C. A. Cobbold, 2012 Measuring diversity: The importance of species similarity. Ecology 93: 477–489.

Lewontin, R. C., 1972 The apportionment of human diversity. In Evolutionary Biology. Springer, 381–398.

Lin, J., 1991 Divergence measures based on the Shannon entropy. IEEE Transactions on Information Theory 37: 145–151.

Louca, S., S. M. S. Jacques, A. P. F. Pires, J. S. Leal, D. S. Srivastava, et al., 2017 High taxonomic variability despite stable functional structure across microbial communities. Nature Ecology and Evolution 1: 1–12.

Louca, S., M. F. Polz, F. Mazel, M. B. Albright, J. A. Huber, et al., 2018 Function and functional redundancy in microbial systems. Nature Ecology and Evolution 2: 936–943.

Lozupone, C., and R. Knight, 2005 UniFrac: A new phylogenetic method for comparing microbial communities. Applied and Environmental Microbiology 71: 8228–8235.

Lozupone, C. A., J. I. Stombaugh, J. I. Gordon, J. K. Jansson, and R. Knight, 2012 Diversity, stability and resilience of the human gut microbiota. Nature 489: 220–230.

McMurdie, P. J., and S. Holmes, 2013 Phyloseq: An R package for reproducible interactive analysis and graphics of microbiome census data. PLoS ONE 8: e61217.

Mehta, R. S., G. S. Abu-Ali, D. A. Drew, J. Lloyd-Price, A. Subramanian, et al., 2018 Stability of the human faecal microbiome in a cohort of adult men. Nature Microbiology 3: 347–355.

Morrison, M. L., N. Alcala, and N. A. Rosenberg, 2022 FSTruct: An FST-based tool for measuring ancestry variation in inference of population structure. Molecular Ecology Resources 22: 2614–2626.

Morrison, M. L., L. Mangé, S. Senkin, N. A. Rosenberg, M. Foll, et al., 2023 Variability of mutational signatures is a footprint of carcinogens. medRxiv.

Nayfach, S., B. Rodriguez-Mueller, N. Garud, and K. S. Pollard, 2016 An integrated metagenomics pipeline for strain profiling reveals novel patterns of bacterial transmission and biogeography. Genome Research 26: 1612–1625.

Oakley, B. B., H. S. Lillehoj, M. H. Kogut, W. K. Kim, J. J. Maurer, et al., 2014 The chicken gastrointestinal microbiome. FEMS Microbiology Letters 360: 100–112.

Oh, J., A. L. Byrd, M. Park, H. H. Kong, and J. A. Segre, 2016 Temporal stability of the human skin microbiome. Cell 165: 854–866.

Olm, M. R., D. Dahan, M. M. Carter, B. D. Merrill, F. B. Yu, et al., 2022 Robust variation in infant gut microbiome assembly across a spectrum of lifestyles. Science 376: 1220–1223.

Paradis, E., and K. Schliep, 2019 Ape 5.0: An environment for modern phylogenetics and evolutionary analyses in R. Bioinformatics 35: 526–528.

Patil, G. P., and C. Taillie, 1982 Diversity as a concept and its measurement. Journal of the American Statistical Association 77: 548–561.

Ricotta, C., 2005 Additive partitioning of Rao’s quadratic diversity: A hierarchical approach. Ecological Modelling 183: 365–371.

Roodgar, M., B. H. Good, N. R. Garud, S. Martis, M. Avula, et al., 2021 Longitudinal linked-read sequencing reveals ecological and evolutionary responses of a human gut microbiome during antibiotic treatment. Genome Research 31: 1433–1446.

Seekatz, A. M., M. K. Schnizlein, M. J. Koenigsknecht, J. R. Baker, W. L. Hasler, et al., 2019 Spatial and temporal analysis of the stomach and small-intestinal microbiota in fasted healthy humans. mSphere 4: 10.1128/msphere.00126–19.

Shalon, D., R. N. Culver, J. A. Grembi, J. Folz, P. V. Treit, et al., 2023 Profiling the human intestinal environment under physiological conditions. Nature 617: 1–11.

Sheth, R. U., M. Li, W. Jiang, P. A. Sims, K. W. Leong, et al., 2019 Spatial metagenomic characterization of microbial biogeography in the gut. Nature Biotechnology 37: 877–883.

Thompson, L. R., J. G. Sanders, D. McDonald, A. Amir, J. Ladau, et al., 2017 A communal catalogue reveals Earth’s multiscale microbial diversity. Nature 551: 457–463.

Tropini, C., K. A. Earle, K. C. Huang, and J. L. Sonnenburg, 2017 The gut microbiome: Connecting spatial organization to function. Cell Host and Microbe 21: 433–442.

Turnbaugh, P. J., R. E. Ley, M. Hamady, C. M. Fraser-Liggett, R. Knight, et al., 2007 The Human Microbiome Project. Nature 449: 804–810.

Upadhyay, V., R. K. Suryawanshi, P. Tasoff, M. McCavitt-Malvido, R. G. Kumar, et al., 2023 Mild SARS-CoV-2 infection results in long-lasting microbiota instability. mBio 14: e0088923.

Xie, F., W. Jin, H. Si, Y. Yuan, Y. Tao, et al., 2021 An integrated gene catalog and over 10,000 metagenome-assembled genomes from the gastrointestinal microbiome of ruminants. Microbiome 9: 1–20.

Xue, K. S., S. J. Walton, D. A. Goldman, M. L. Morrison, A. J. Verster, et al., 2023 Prolonged delays in human microbiota transmission after a controlled antibiotic perturbation. bioRxiv : 2023.09.26.559480.

Yassour, M., T. Vatanen, H. Siljander, A.M. Hämäläinen, T. Härkönen, et al., 2016 Natural history of the infant gut microbiome and impact of antibiotic treatment on bacterial strain diversity and stability. Science Translational Medicine 8: 343ra81.

Zaneveld, J. R., R. McMinds, and R. V. Thurber, 2017 Stress and stability: Applying the Anna Karenina principle to animal microbiomes. Nature Microbiology 2: 1–8.

